# The role of hippocampus during observational learning

**DOI:** 10.1101/832758

**Authors:** Yerko Fuentealba, José L Valdés

**Author notes:** Corresponding author: José L Valdés, Departamento de Neurociencias, Facultad de Medicina, Universidad de Chile. Independencia 1027, Independencia, Santiago, Chile.

## Abstract

Observational learning is a fundamental cognitive ability present in several species, where a naïve animal imitates a goal-directed behavior from the observation of a congener which acts as a demonstrator. Recent evidence in bat and rats suggests that hippocampal place cells of an observer may generate a spatial representation of the locations visited by a demonstrator, during spatial navigation. However, it is still unclear whether this hippocampal neural activity is critical for the process of observational learning or if the patterns of activity during observation differ from those emerging from the execution of a spatial memory task previously observed. To test this idea, we assess the role of the hippocampus by pharmacological reversible inactivation during the observation of a spatial learning task, demonstrating a critical role for this structure in observational learning. Then we recorded the neuronal activity of principal pyramidal cells of the same animal when it was observing or solving the memory task, and two different representation of the space emerged after observation or navigation. This evidences demonstrated that the hippocampus is necessary for observational learning and indicated that the observed and executed hippocampal representation are different, confirming the idea that the hippocampus could represent the position of others in the space, and use this information to improve his behavioral performance.

## INTRODUCTION

Observational learning is the ability of an individual to imitate a goal-directed behavior from the observation of a congener that acts as a demonstrator. This ability ensures the transference of adaptative behaviors through generations in different species (Bandura 1969, Zentall 1996, Heyes and Galef, 1996, Galef 1988, Marler and Tamura, 1964; Fiorito and Scotto, 1992). In rodents, observational learning has been demonstrated in operant conditioning (Heyes and Dawson, 1990), fear conditioning (Jeon et al., 2006) and spatial navigation tasks (Leggio et al., 2000). A characteristic feature of this learning is that the observer does not necessarily repeat the behavior of the demonstrator in a movement-to-movement way. Observers develop their own repertoire of movements to achieve the same goal that the demonstrator, suggesting that the observer understands the meaning or aim of the behavior displayed by the demonstrator (Tomasello M., 1996). A neurophysiological substrate for this cognitive ability could be the generation of internal representation or cognitive map (Tolman, 1948) in the brain of the observer concerning the demonstrator behavior.

Place cells in the hippocampus provide a neurophysiological framework for a cognitive map, which emerges during spatial navigation and underlies the learning and memory process (O’Keefe and Nadel, 1978). Place cells increase their firing rate in specific locations (place field) of the environment traveled by the animal (O’Keefe and Dostrovski, 1971), and subsequently, multiple place fields generate a complete representation of the environment explored by an animal, the spatial or cognitive map (O’Keefe and Nadel, 1978, Barnes et al., 1997). This hippocampal cognitive map is flexible enough to generate multiple representations of multiple experiences of the animal, through the remapping phenomenon (Leutgeb et al., 2005). The generation of a cognitive map has been widely described for rodent navigation, and recent evidence has indicated that hippocampal CA1 cells of an observer animal may code for the position of a demonstrator (Denjo et al. 2018, Omer et al. 2018). Trained rats and bats were challenged to choose a left or right path in a two-alternative maze based on the direction taken by a demonstrator. Electrophysiological recording during the observation of this task showed that a subset of neurons (social place cells) encodes the position of the demonstrator. Even though this evidence strongly suggests that the hippocampus may be relevant for the establishment of observational learning, a causal relationship between the hippocampus and the capacity to learning by observation has not been clearly explored.

In the present study, we developed an observational learning spatial memory task, where naïve animals, without any experience in the task, observed a well-trained demonstrator, to determine the role of the hippocampus in observational learning. After reversible inactivation of the hippocampus during observation, we establish a causal relationship between hippocampal activity and observational learning. By using high-density electrophysiology in the behaving rats, we determine the main differences in the activity of social place cells during observation and typical place cells during task execution.

## METHODS

### Subjects

Fifty-eight adult male *Sprague-Dawley* rats were used. All animals were obtained from our institutional Animal Care Facility; weighing 270-320g. They were individually housed with *ad libitum* access to water and food, at least the other was indicated, in a temperate room (23°C) and light/dark cycles of 12/12 hrs, ZT0=7:00 AM. Surgical and experimental procedures were carried out in accordance with the National Institute of Health (USA) Guide for the Care and Use of Laboratory Animals (NIH Publications No. 80-23, revised 1996). The institutional Biosafety and Ethical Committee (CBA# 0770, FMUCH, University of Chile) approved these experimental protocols, which minimized the number of rats used and their suffering.

### Animal groups

For behavioral experiments, we used 33 animals that were divided into two groups as follow:

Group 1, D*emonstrators:* 14 rats were used as a demonstrator of a spatial learning task. Seven of them were highly experimented demonstrators who were pretrained to solve the task 3 days before the test (pretrained group). The other seven rats were also demonstrators but without any experience or pretraining to solve the maze (Naïve group).

Group 2, Observers: 19 rats were used as observers and were initially habituated in the observational platform of the maze Fig. 1A, (see section *Spatial learning task* for details) during three consecutive days. The fourth day this observer animal group were divided on those that watched a pretrained demonstrators (observers of a pretrained group, n=6), and those observers that watched a naïve demonstrator (observers of a naïve group, n=5) and the remaining observers did not watch any demonstrator (control group, n=8). Comprehensive animal group details are indicated in Fig. 1B and Fig. S1.

**Figure 1.**
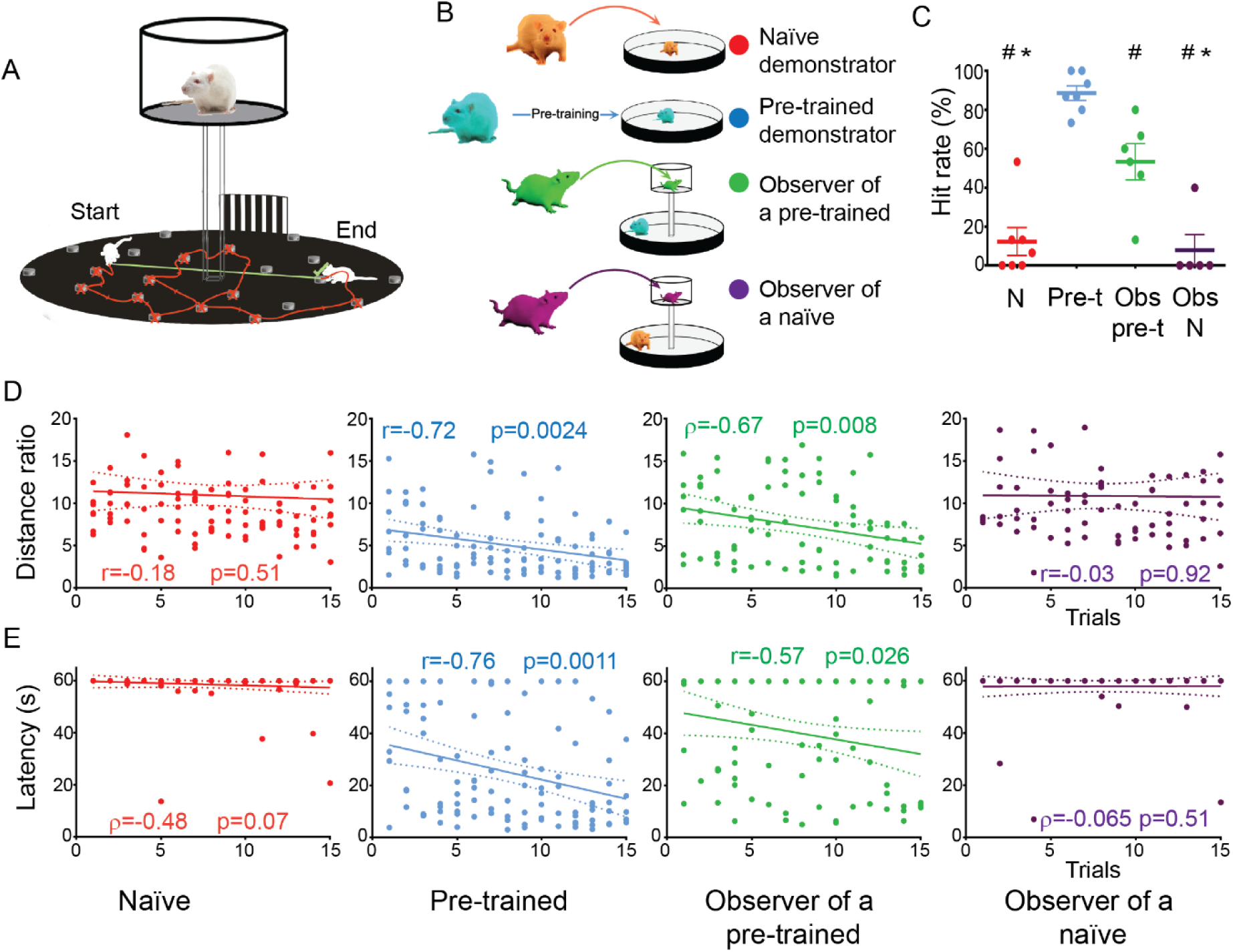
Observation improves spatial learning in the oasis maze. **(A).** A modified version of the Oasis maze with an observation platform on it where a naïve rat observed a demonstrator rat solving the task. The green line depicts the optimal path between the start and the end while the red one indicates a path traveled by the rat. **(B).** Behavioral groups: rats were divided into demonstrators (red and blue) and observers (green and purple). The demonstrators were separated into naïve and pre-trained animals, while the observers were always naïve and separated based on the demonstrator: observer of a pre-trained (green) or an observer of a naïve (purple). **(C).** Mean ± sem percentage of hit rate. of four experimental groups: not pre-trained (“N”, red), pre-trained (“Pre-t”, cyan), an observer of a pre-trained (“Obs pre-t”, green), an observer of a not pre-trained (“Obs N”, purple). #p<0.05 vs pre-trained group; *p<0.05 vs observer of a pre-trained group; one-way ANOVA, followed by Holm-Sidak posthoc test. **(D).** Distance ratio: the ratio between the actually walked distance over the optimal distance, on each trial. Each plot represents one of the four experimental groups described before. **(E).** Latency: time spent for rats to find the rewarded well on each trial. In (D) and (E) the correlation coefficient and p-value of each group are indicated, (Pearson correlation coefficient or Spearman Rank correlation index).

Twenty-five additional animals undergo surgical procedures, 18 animals were bilaterally implanted with injections cannulas targeting the hippocampal dorsal CA1 region, and seven rats were implanted with a seven tetrodes-hyperdrive array for *in vivo* electrophysiological recording of CA1 neuronal activity.

### Spatial learning task

To test spatial learning we used a modified version of the oasis maze, a dry land version of the Morris water maze (Clark et al., 2005, Martinez et al., 2016). The apparatus consisted of an open field arena of 140 cm in diameter, elevated 50 cm from the floor, with a wall of 20 cm of height. Twenty equidistantly evenly spaced wells were distributed over the arena. The rats were water-deprived for 23 hours to enhance motivation and pretrained to seek for water (200 µl water drop) inside of the wells during three consecutive days, up to the animals were able to find the 100% of baited wells for 15-20 minutes session per day. Twenty, fifteen and five wells were daily baited during the pretrained period. After the pretraining, the task consisted of 15 trials of 1 minute each with only one baited well, preserving the position of the well along with the task. The starting position of the animal was changed on each trial to avoid the development of stereotyped procedural behavior, and after each trial the rat was enclosed with a black carboard-cylinder of 22 cm in diameter and 27 cm in height over the arena and gently moved to a new starting position, with the aim of preventing handling during task execution. The trial started after the cylinder was removed and ended when the rat reaches the rewarded well or one minute elapsed, 20 to 30 seconds of the inter-trial interval was included, where the animal was gently moved to a new starting position Fig. 1A.

### Observational learning task

An elevated platform (50 cm of the floor, 35 cm in diameter and 40 cm of height walls) built on transparent polymethacrylate and a grid mesh floor, was in the center of the oasis maze arena (Figure 1A). The observer rats were placed in the elevated platform while a demonstrator solves the task in the oasis arena. Thirty consecutive trials of observation were conducted. Immediately after the observation, the observer rats were challenged to solve the maze.

### Pharmacological inactivation of the hippocampus

*O*bserver animals were chronically implanted with bilateral drug infusion cannulas (23 ga guide cannula, 33 ga injection cannula, Plastics One, Inc, VA, USA), targeting the dorsal CA1 region of the hippocampus to reversibly inactivate the hippocampal neuronal activity. One injection of 0.5 µl of bupivacaine (0.75% v/v, Abbott Laboratories, IL, USA) or vehicle (saline, NaCl 0,9%) was infused by using a 10 µl Hamilton syringe at a rate flow of 0.5 µl/min. After injection, we leave the cannula in the site for one additional minute to allow proper drug diffusion, experimental animal groups detailed in Fig S2.

**Figure 2.**
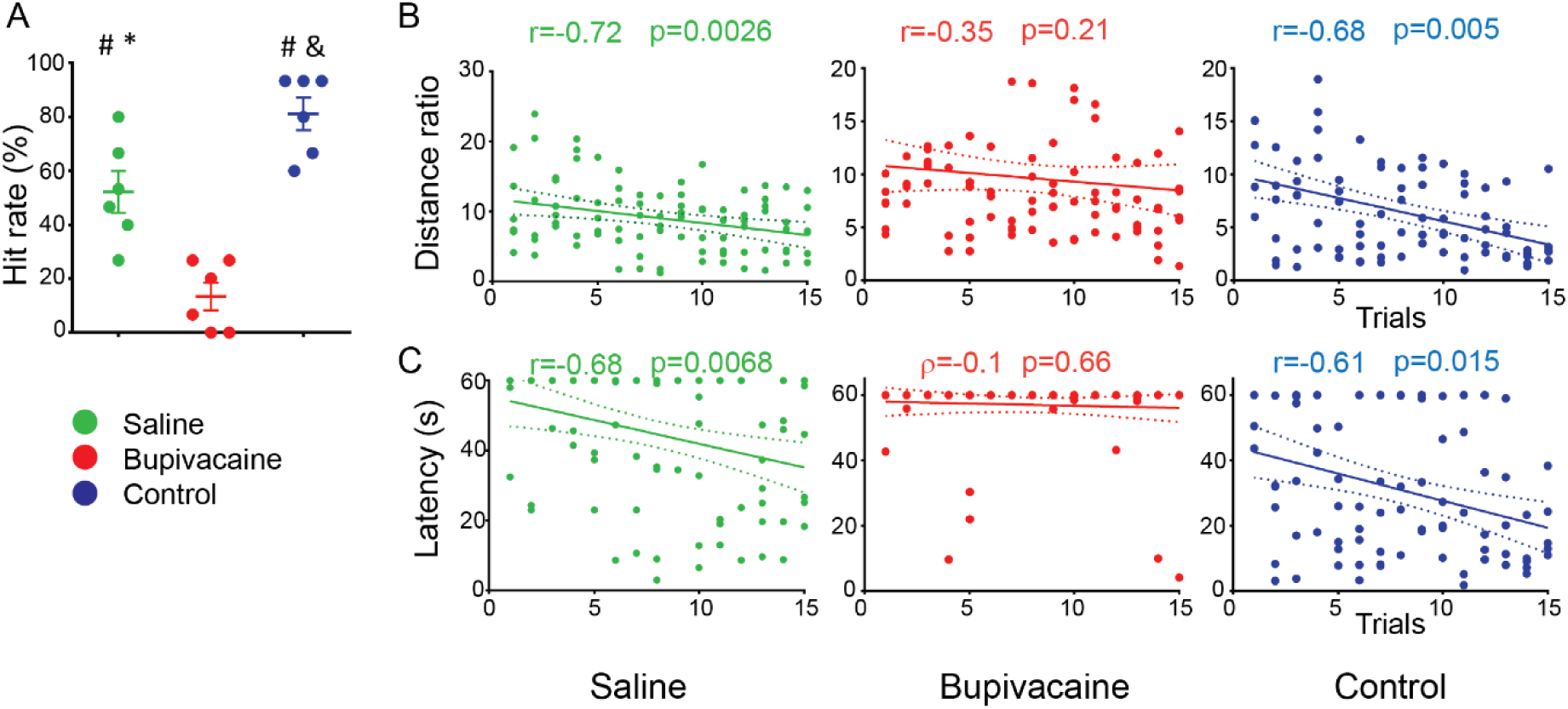
Pharmacological inactivation of the hippocampus during observation abolishes spatial learning. Rats were separated into three groups based on the injection: saline (green), bupivacaine (red) or control (blue). Control animals, previously pretrained in the oasis maze, were injected with bupivacaine 40 minutes before the task. **(A).** Mean±sem percentage of hit rate. & p<0.01 vs saline group, #p<0.01 vs bupivacaine group, *p<0.01 vs control (one-way ANOVA followed by Holm-Sidak posthoc test). **(B)** distance ratio, and **(C)** latency for each group. The correlation coefficient and p-value of each group are indicated, (Pearson correlation coefficient or Spearman index).

### Surgical procedures

All rats implanted with cannulas or electrophysiological recording electrodes were anesthetized with 2.5% of Isoflurane/Oxygen mixture gas for induction and 1.5% for maintenance at a flow rate of 1 l/min. Antibiotic (Enrofloxacin, 19 mg⁄kg, i.p.; Bayer) and anti-inflammatory (Ketophen 0.2 mg⁄kg, i.p.; Rhodia Merieux) were administered at the end of surgery and during three additional days. The animal was fixed in a stereotaxic frame, a small incision in the scalp and a craniotomy was made. The electrodes or cannulas were implanted through the craniotomy. All implants were fixed to the skull with five anchor jewelry stainless steel screws and dental acrylic. One of the skull screws was used as an animal ground for electrophysiological recording. Eighteen rats were implanted with cannula guide at 3.3 mm posterior to bregma; ± 2.5 mm lateral, with a 20° angle respect to the midline and 1.2 mm in dept, following the Paxinos’s rat atlas coordinates (Paxinos and Watson 1998). The injection cannula was 1mm longer than the cannula guide. Seven rats were chronically implanted with a hyperdrive consisting of six independent movable tetrodes. The six tetrodes were targeted to the dorsal CA1 region at 3.3 mm posterior to bregma, 1.8 mm lateral to the midline and ∼2.2mm in dept, following the Paxinos’s rat atlas coordinates (Paxinos and Watson 1998).

### Electrophysiological data acquisition and data analysis

Electrophysiological signals were recorded from each of the 24 wires (six tetrodes) simultaneously. The tetrodes (Wilson and McNaughton 1993) were made by twisting four 17 µm nichrome insulated wires (AM systems, USA), gold plated to the impedance of 0.5-1 MΩ. Each tetrode was independently lowered to the target area, at a rate of no more than 320 µm/day, until appropriate signals could be recorded. The leads of the tetrodes were connected to a unity-gain headstage, and all the data were collected using analog-32 Channels, Cheetah recording System (Neuralynx, Bozeman, MA, USA). Single unit data from each tetrode was amplified, bandpass filtered (600–6,000 Hz) and digitized at a rate of 32 kHz. LFP signals were acquired with the same system, filtered between 0.1-450 Hz and digitized at a rate of 2 kHz. Single neurons were isolated offline using automatic cell sorting software KlustaKwik (by K. Harris) and manually supervised with the software MClust (by D. Redish). All spikes clusters with more than 1% of interspike intervals lower than 2 ms were considered as multi-units and discarded for further analysis (Valdes et al., 2015). The animal behavior was video recording simultaneously with the electrophysiological recording.

### Hippocampus cell classification and place cell determination

We obtained 380 neurons from 7 rats in 10 experimental sessions. Putative pyramidal and interneurons were classified in agree with their firing rate and waveform: units with a firing rate lower than 0.25 Hz during observation or navigation were discarded since its low firing rate does not support further analysis (n=41). Neurons with a firing rate higher than 10 Hz and a peak-to-through waveform ratio close to 1 were considered putative interneurons and discarded for further analysis (n=17), all the rest of neurons (n=322) were classified as potential pyramidal cells.

We build rate maps for all pyramidal neurons by dividing the oasis maze arena into 48×48 bins (∼3.2 x 4.2 cm each bin) and computing the instantaneous firing rate on each bin (spikes per occupancy). During the observation phase of the task, we used the occupancy of the demonstrator rat, and for the navigation phase the occupancy of the observer while it was solving the maze. The firing rate of each neuron was normalized as follow:

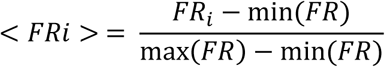

With *FRi*= Firing rate at bin (i) and maximum and min of the firing rate (*FR*) during the analyzed epochs. Then, the information per spike index (IPS) of each neuron during observation and navigation was calculated based on (Skaggs et al., 1993; Robbe and Buzsaki, 2009):

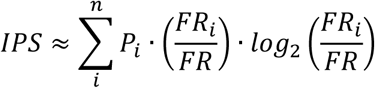

Where *i* is the bin number, P_i_ is the probability of occupancy at the bin *i*, FR_i_ is the firing rate in the bin _i_ and FR is the overall mean firing rate. The IPS computing was using the non-normalized values of FR_i_. The IPS calculated for each pyramidal neuron was compared with a randomized IPS, that was calculated by bootstrapping the spiking time of each spike train time series, preserving the interspike interval (ISI). This procedure was conducted 1,000 times, then all those IPS higher than the mean ± 2SD of the randomized IPS were considered as pyramidal cells with significant spatial information content over the chance. Those cells which have an IPS higher than chance during observation were classified as social place cells, those neurons with a significant IPS during navigation were classified as canonical place cells, those cells which support the previous criteria on both phases of the task (observation and navigation) were classified as common cells.

Sparsity was calculated as (Jung et al., 1994):

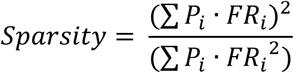

Where *P_i_* is the probability of occupancy of bin *i* and *FR_i_* is the firing rate in the bin *i. Spatial correlation*. Rate maps for those neurons classified as social place cells and place cells for the same experimental session were compared by vectorizing the 48×48 rate maps and computing the bin-by-bin Pearson correlation index between observation and navigation phases (Leutgeb et al., 2005). We discarded all those bins that in both phases showed no activity. The spatial correlation of each neuron was compared with the spatial correlation of the observation rate map and the navigation rate map of other neurons (spatial correlation by chance). Then spatial correlation lower than 2SD of the spatial correlation computed by chance indicates completely unrelated rate maps (Fyhn et al., 2007).

### Place field firing rate overlap

Differences in maximum firing rates between the phase of observation and navigation were evaluated by computing the overlap score of the firing rates for each neuron classified as social place cell and canonical place cells. Overlap score for each neuron was calculated as the ratio between higher maximum firing rate of observation or navigation phases of the task, to get values in the interval of 0 to1, where 1 indicates no difference and values closer to 0 indicates a greater difference between the two phases of the task (Leutgeb et al., 2005). To statistically determine rate overlap (rate remapping), we computed the overlap score between each common cell during observation and all other common cells during navigation to estimate the overlap by chance, then we consider rate remapping when the overlap score is higher than 2SD of the overlap computed by chance.

### Population vector analysis

A population vector analysis was performed (Leutgeb et al., 2005, Roux et al., 2017) to evaluate the similarity between observation and navigation representation at the neural population level. To this, a stack of the firing rates of each bin of all common cells of a same experimental session was adjusted in a 3D matrix, where the x- and y-axis represents the position in the maze and the z-axis corresponded to the firing rate of each neuron, in both phases of the task (observation and navigation). For each bin, the population vector (PV) corresponded to the distribution of normalized firing rates of different neurons in the same bin. Then, we calculated the Pearson (bin-by-bin) correlation index for each PV between observation and navigation and compared this distribution of PV correlation with a random-generated distribution obtained after shuffling the position of different neurons in the z-axis of each bin, for each experimental phase. This randomization was iterated 1000 times, and the distributions of raw and random-generated PV correlations were compared.

### Theta band power spectral density analysis

The theta band (4-12 Hz) power spectral density was calculated using the fast Fourier Transform with the multitaper method (Mithra and Bokil, 2007). The tetrode with the best LFP signal (highest amplitude and low noise) was selected for the analysis. Raw power spectral density values were normalized by its maximum and minimum values for statistical comparison among task phases, experimental sessions, and animals. A spectrogram in the 0.1-20 Hz band built with the multitaper method using chronux toolbox in MATLAB (http://chronux.org/, Mithra and Bokil, 2007) with K=5 and TW=3, without subsampling the continuous signal.

### Statistics

All statistical analysis was performed using the software GraphPad Sigma 6.01. All results were expressed as mean ± s.e.m. The electrophysiological analysis was performed by using custom-made MATLAB (Mathworks, Inc) routine. In all cases, the normality of the data was assessed with the Kolmogorov test before to compute any statistical comparison. Then, we used non-parametric tests when the data did not fit the normal distribution. The statistical significance was fitted to a p-value <0.05.

The comparison of different behavioral variables was mainly performed by using one-way ANOVA followed by a multiple comparisons Holm-Sidak posthoc test, or Kruskal Wallis test followed by Dunn’s posthoc. The correlation between distance ratio and latency and the progression of the task (‘trials’) was evaluated by Pearson correlation index or Spearman rank-correlation coefficient.

The comparison among mean/max firing rate, information per spike index and sparsity was performed using a Mann-Whitney rank-sum test. For firing rate maps comparison, the Pearson correlation index bin-by-bin was used. For the max firing rate comparison in the same neuron during different epochs of the task (common neurons), we used the Wilcoxon paired rank-sum test. Finally, the theta range (4-12Hz) power spectral density in rest, observation and navigation, were compared using the Friedman test followed by Dunn’s comparison.

## RESULTS

### Naïve rats improve their performance in a spatial navigation task trough observation

To evaluate the ability of rats to learn by observation, we implemented a modified version of the spatial learning task (Oasis maze), where naïve animals can observe a demonstrator and then execute the same task (Fig. 1A-B). We used three parameters to evaluate spatial learning performance: hit rate (percentage of successful trials, Fig. 1C); distance ratio (ratio between traveled distance and the optimal distance) and latency (time spent by the animals to get the rewarded well). Distance ratio and latency were used to evaluate spatial learning as a progressive decrease of each variable through the trials. The Naïve animals (Fig 1C, “N” in red) showed the lowest hit rate and not progression across trials for distance ratio (Fig. 1D, in red, r=−0.18, p=0.51) or latency (Fig 1E, in red, ρ=−0.48, p=0.07), in the oasis maze. In contrast the pretrained animals (cyan on Fig 1C-E) showed the highest hit rate (Fig. 1C, “Pre-t”, in cyan) and proper progression of learning through trials for distance ratio (Fig 1D, cyan, r=−0.72, p=0.0024) and latency (Fig 1E, cyan, r=−0.76, p=0.0011). Those animals which observed a pretrained demonstrator (observer of a pretrained, Fig 1C-E in green), showed a higher hit rate (Fig 1C, “Obs pre-t”, in green) and significant progression in distance ratio (Fid 1D, green ρ=−0.67, p=0.008) and latency (Fid 1E, green, r=−0.57, p=0.026) while those animals which observe a not pretrained animal (observer of a not-pretrained, Fig 1C-E, in black) did not show any significant improvement in their spatial learning performance with a low hit rate (Fig 1C, “Obs N”, in black) and no progression in distance ratio (Fig. 1D, in black, r=−0.029, p=0.92) or latency (Fig. 1E, in black, ρ =−0.07, p=0.5). As an additional control, rats only habituated to the platform without any observation of a demonstrator showed no significant learning performance in all the variables previously indicated (Fig. S3).

To rule out the possibility that poor spatial learning could be explained by low displacement in the arena or even immobility, we analyzed the total path length traveled through trials and the mean velocity of each animal for each group (Fig S4). Pretrained rats showed the lower total path length (61.82±9.73 cm) and the highest velocity (18.39±0.52 cm/sec) compared with rats that did not show progress in learning (Not pretrained, 96.28±6 m and 11.06±0.28 cm/sec; Observer of a not pretrained, 97.82±4 m and 11.54±0.41 cm/sec, Fig. S4). This result indicates that those animals which did not properly solve the task explored the arena even more than those which solved it. Then, the absence of learning could not be explained due to immobility or absence of exploration in the maze. These results indicate that only those animals which have the chance to observe an experimented pretrained demonstrator improve their learning performance. Conversely, those animals which watched a naïve demonstrator, solving the task for the first time or they only were habituated to the platform, display poor performance in the oasis maze. The observational learning could not be attributable to the mere presence of a congener in the arena or the habituation to the observational platform.

### Hippocampal inactivation abolishes observational learning

To determine whether the hippocampal activity is necessary for observational learning, we pharmacologically inactivated the dorsal CA1 region during observation and evaluated its impact on observational learning performance. Animals were bilaterally injected with 0.5 µl of bupivacaine 0.75% v/v. This sodium channel blocker was used since it has an *in vivo* half-life about 40 minutes (Catterall and Mackie, 2012), which matches with the duration of the observation phase of our task, allowing us to differentiate the requirement of hippocampal activity exclusively during observation, without affecting the functionality of this structure in the subsequent phase of spatial learning during the task execution.

Two groups of observer rats were injected with bupivacaine, or saline during the period of observation. Immediately after drug injection, both groups observed the behavior of a pretrained demonstrator in the same conditions as before. To discard any remnant effect of bupivacaine in the functionality of hippocampus during the testing period of navigation, we included a third control group of previously pretrained animals in the oasis maze and injected with bupivacaine 40 minutes before solving the task (control), but without observation (detail of animal groups in Fig. S2).

Observer rats injected with saline showed 52.22 ± 7.78 % of hit rate, which was higher than the bupivacaine-injected animals which showed a 13.33 ± 5.16 % of hit rate (Fig. 2A, p=0.0013). Control-injected animals, in turn, showed the highest hit rate among all groups (Fig. 2A, 81.11 ± 6.07 %, p=0.0062 vs. saline and bupivacaine p<0.0001, one-way ANOVA on ranks, followed by Holm-Sidak multiple comparisons test). Saline-injected animals showed a decrease in distance ratio and latency along the task (r=−0.72, p=0.0026 and r=−0.68, p=0.005 respectively), that was not found in bupivacaine-injected animals (r=−0.35, p=0.21 and ρ=−0.1, p=0.66 respectively), control-injected animals showed normal decrease in distance ratio and latency (r=−0.68, p=0.005 and r=−0.61, p=0.015 respectively, Fig. 2B-C).

The results obtained for the hit rate after pharmacological inactivation of the hippocampus resembled the results obtained in the previous experiment: saline-injected animals reaching the same hit rate than observers of a pretrained demonstrator and displayed a significant decrease in both variables distance ratio and latency. Bupivacaine-injected animals showed a similar hit rate of naïve and observers of a not-pretrained, without a significant progressive decrease in distance ratio and latency. Finally, the control-injected group reached the same hit rate as pretrained rats and decreased their distance ratio and latency along with the task (Fig. S5). These results indicate that hippocampal function is essential for observational learning since its inactivation prevents learning improvement. The absence of learning could not be attributable to the remnant effect of the drug during spatial learning, because pretrained animals injected with bupivacaine 40 minutes before the test, solving the maze with high proficiency, indicating that after that time the hippocampal function was fully recovered.

### The emergence of social place cell activity

Social place cells have been recently described as CA1 hippocampal pyramidal neurons that can generate a representation of a congener in socially guided tasks, both in rats and bats (Danjo et al., 2018, Omer et al., 2018, Bray, 2018, Duvelle and Jeffery 2018). We tested the idea that social place cell-activity could underlie observational learning during a spatial learning task. Given that observer rats were always naïve, this hypothesis also implies that observers could generate a spatial representation of unvisited environments, by representing the positions of the demonstrator. To test this idea, we recorded the neuronal activity of CA1 hippocampal cells of a naïve observer while a pretrained demonstrator performs the spatial learning task. After recording, single units of putative pyramidal neurons were analyzed following the firing rate and spike waveforms criteria defined in the methods section. A total of 380 individual neurons from 7 animals in 10 experimental sessions were obtained, 85% of which were classified as putative pyramidal cells (n=322 neurons). Low firing rate neurons or putative interneurons were not included in these analyses. To determine whether a pyramidal neuron coded for the position of the demonstrator we aligned all the trajectories traveled by the demonstrator rats with the timestamps of each spike for each pyramidal neuron of the observer hippocampus. Then we built rate maps for each neuron as was described before.

For each rate map, we computed the information per spike index (IPS, Skaggs et al., 1996) and compared this value with the IPS obtained by chance (1,000 randomizations of the interspike interval of each rate maps (Omer et al., 2018). All those neurons that showed an IPS higher than 2SD over the mean of the randomized IPS were classified as social place cells since its information content was higher than expected by chance. We identified 19,3% (62 neurons) of total pyramidal neurons as “social place cells” during the observation phase of the task. These neurons will be encoding the position of the demonstrator in the observer’s hippocampus (Fig 3A). By using the same criteria, we classified 26.4% (85 neurons), of pyramidal neurons as a place cells during the navigation phase of the task (Fig. 3B).

**Figure 3.**
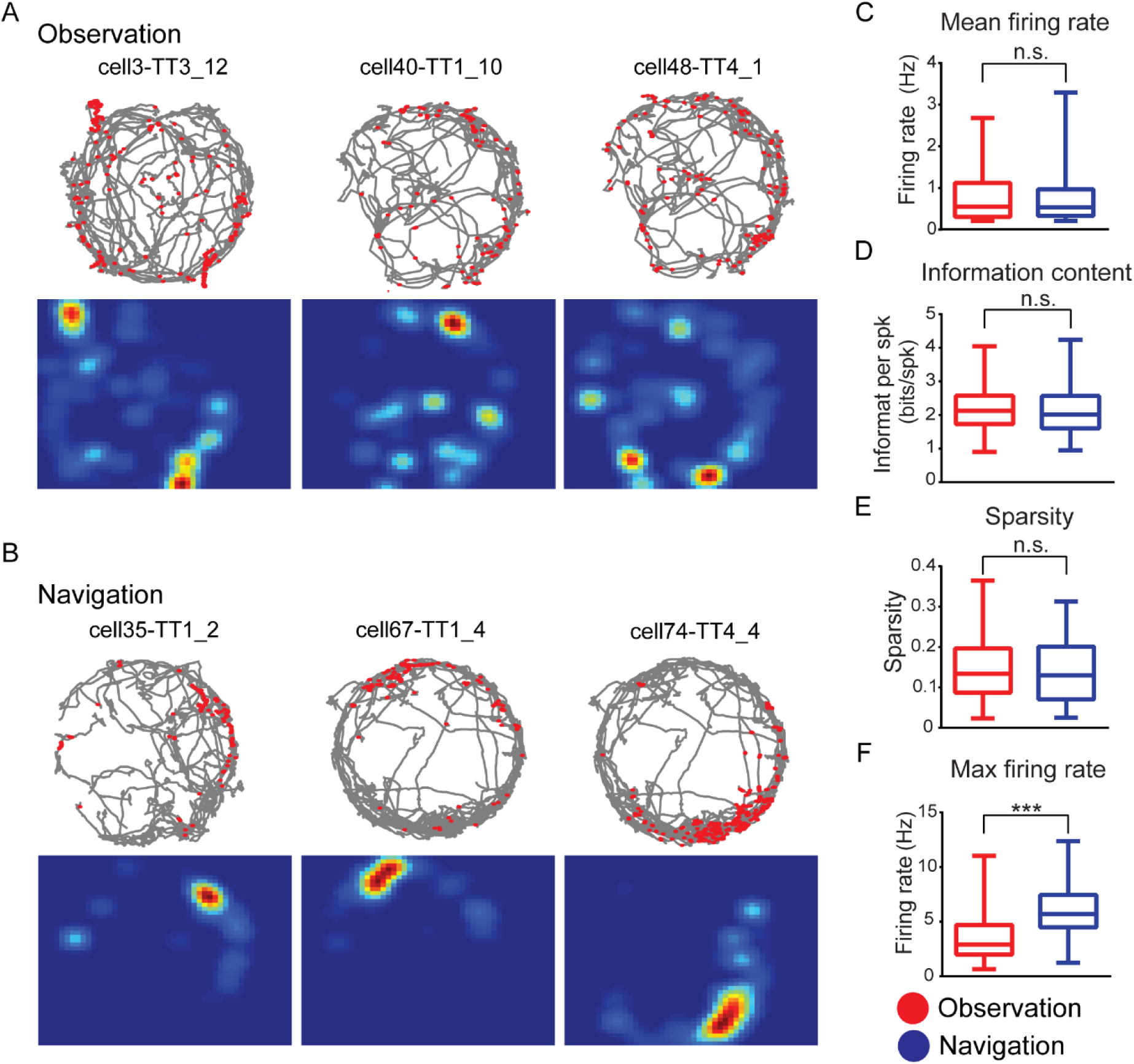
Social place cell activity during observational learning. **(A)** Representative examples of observer’s hippocampal CA1 neuronal activity (upper panel, animal paths in grey lines and neuron spikes in red dot) with respect to demonstrator trajectories (lower panel, rate maps with normalized firing rate). **(B)** Representative examples of observer’s hippocampal CA1 neuronal activity during navigation. **(C)** Mean firing rate during observation (red) or navigation (blue). **(D)** information per spike (IPS) during observation or navigation. **(E)** sparsity and **(F)** maximum firing rate (at the center of the place field) for the population of neurons active during observation (red) and navigation (blue). ***p<0.001 Mann-Whitney rank-sum test.

Social place cells obtained during the observation phase shared several features with canonical place cells recorded during navigation, with no statistical differences in its mean firing rate, information per spike content and sparsity of the place fields between the two phases of the task (Fig. 3C-E). Only the maximum firing rate (at the center of the place field) was lower during observation than during navigation (3.64±0.3 Hz and 6.13±0.26 Hz respectively, p<0.0001 Mann-Whitney rank-sum test, Fig. 3F).

These results suggest that during observation, a subpopulation of CA1 pyramidal neurons can encode the position of the demonstrator in the hippocampus of the observer. These neurons shared several features of canonical place cells, such as the mean firing rate, the information per spike index and sparsity, suggesting that the social place cells and place cells are the same neurons. However other properties, as the maximum firing rate in the center of the place field was lower during observation that during navigation, suggesting that the cognitive maps emerging from observation and navigation are different at least regarding firing rate.

### The spatial representation during observation is not correlated with spatial representation emerging from the navigation

To determine a relationship between the cognitive maps generated during observation and navigation, we focused on the fraction of neurons that showed place cell activity during both phases of the task, that we called common cells. We identified 29 common cells from the 62 observational place cells or 85 place cells (46.8% and 34.5%, respectively, Fig. 4A). This data indicates that almost half of social place cells take part in the representation of the self-animal as place cells, suggesting a significant contribution to the spatial learning behavior. To evaluate whether the spatial map codification of the demonstrator resembles the representation of the self-animal during navigation, we compared the spatial position of the place fields and the firing rate patterns, for each common cell during observation and navigation. First, we compared the spatial representation in both conditions of each common cell by using Pearson bin-by-bin spatial correlation index, Fig. 4B. To determine whether a spatial correlation was higher than chance, we compared the spatial correlation obtained for each cell against a threshold computed as 2SD above the average correlation of completely unrelated maps, which denotes full global remapping (see methods for details, Fhyn et al., 2007). We found that only one out of 29 common cells showed a spatial correlation higher than the expected by chance (Fig. 4C), which indicates that the spatial representation generated by a given common cell during observation is unrelated with the spatial representation generated by the same cell during navigation.

**Figure 4.**
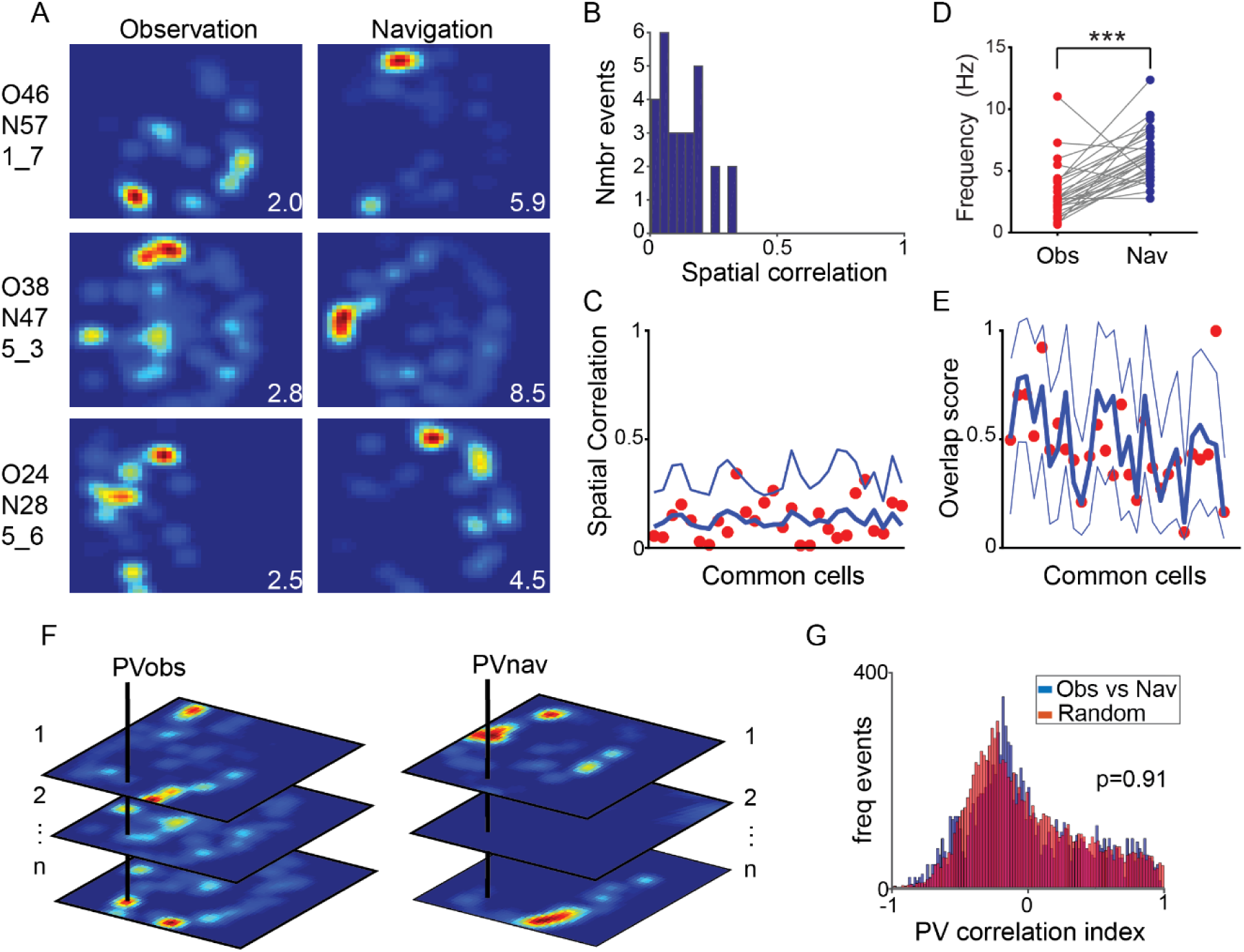
Global remapping during the transition between observation and navigation. **(A)** Examples of the common cells, neurons that were active during observation and navigation. The activity was normalized to the maximum firing rate of each rate map. **(B)** Distribution of spatial correlation index calculates for the same common cell between observation and navigation phases of the task. **(C)** Spatial correlation index obtained for each common cell (red dots). Blue lines indicate the randomized spatial correlation intervals values (mean+2SD, thick and thin lines respectively), neurons outside this interval are considered with similar rate maps, otherwise, the maps are significantly uncorrelated. **(D)** Maximum firing rate of common cells during observation (red) and navigation (blue). ***p<0.0001, Wilcoxon-rank test. **(E)** Overlap score of firing rates during observation and navigation for each common cell. Values close to 1 indicate no rate remapping and values close to 0 the opposite. As in (C), thick blue lines denote the mean random overlap scores and thin blue lines denote the mean ± 2SD random overlap scores. **(F**) Population Vector analysis: The PV of each bin was obtained staking the firing rate of each bin, of the rate map, for each common cell, during observation (PVobs) and navigation (PVnav). **(G).** The comparison of the PV correlation index between observation and navigation was not different than a randomized PV population built by using the same data set (p=0.91, Mann-Whitney test).

Second, we evaluated rate remapping, as a change in the maximum firing rate during the observation and navigation phases of the task. We found that the maximum firing rate of common cells during the observation was lower than navigation (Fig. 4D, p<0.0001, Wilcoxon paired signed-rank). The overlap firing rate score for each common cell indicates that only two out of 29 common cells showed overlap scores higher than the chance, while most neurons (93.1%) showed overlap scores into or below the stablish by chance, suggesting that almost all those neurons exhibit rate remapping (Fig 4E).

Third, we performed a population vector analysis to assess the similarity of the representation at the population level during observation and navigation (Leutgeb et al., 2005). The firing maps of common cells were stacked in a 3D matrix where *x*- and the *y*-axis represents the position of each bin of the map, and the *z*-axis represents the firing rate of each neuron. The population vector (PV) corresponded to the distribution in each bin of normalized firing rates from different common neurons in a single experimental session. We calculated the Pearson correlation bin-by-bin for each PV between observation and navigation and compared this distribution with a randomly generated-distribution (see details in methods). The distribution of PV-correlation coefficients was not different than the random distribution (0.33±0.3.3+10^−3^ and 0.32±3.2*10^−3^ respectively, p=0.91, Mann-Whitney rank-sum test), indicating that there was no correlation at population level higher than the obtained by chance and that the organization of the cognitive maps, at the population level, was different in the two phases of the task (Fig. 4F, G).

Finally, we evaluated whether the cognitive maps generated during observation and navigation preferentially encoded a salient feature of the environment, such as the location of the rewarded-well. The distance (in cm) between the center of the place field and the rewarded well during both phases were computed. We found that the distribution of distances statistically fits with a normal distribution with a mean of 67.48±3.12 (cm) for observation, and 68.79±2.99 (cm) for navigation, that was not different among them (Fig. S6, p=0.76, t-test). This result indicates that place fields were evenly distributed in the oasis maze arena and unbiased by the rewarded well both during observation and navigation phases.

In summary, we found a population of hippocampal neurons that displayed a significant amount of spatial information about the demonstrator’s position during the observation phase. Near half of those neurons were also part of the population of canonical place cells emerging during the navigation phase. Our analysis indicates that the spatial representations during observation and current navigation are entirely different.

### Theta oscillation during observation and navigation

The changes in the LFP signal in the range of theta (4-12 Hz) are typically displayed during spatial navigation, strong temporal coordination between spike cell activity and theta oscillation is present and necessary for learning process (Robbe et al 2006, Skaggs et al 1996) and have been described for social place cells activity (Denjo et al., 2018). To determine whether during observation some similar LFP/spike coupling is also present, we compared changes in theta power during a 10-minute resting period before and after observation or navigation. We did not find differences in power spectral density on the theta band during observation compared with a pre-task resting period (0.46±0.07, dB and 0.48±0.08, dB; respectively, p>0.99, Friedman test followed by Dunn’s test), but during navigation we found an increase in theta spectral power compared with rest (0.63±0.07, dB, p=0.04, Friedman test followed by Dunn’s test, Fig. 5A, B). This data suggests that neuronal activity emerging during observation did not correlate with an increase in the spectral power of the internally-generated theta oscillation, that it is present during navigation, indicating that animals without experience in the observed environment do not coordinate observational place cells activity with theta rhythm.

**Figure 5.**
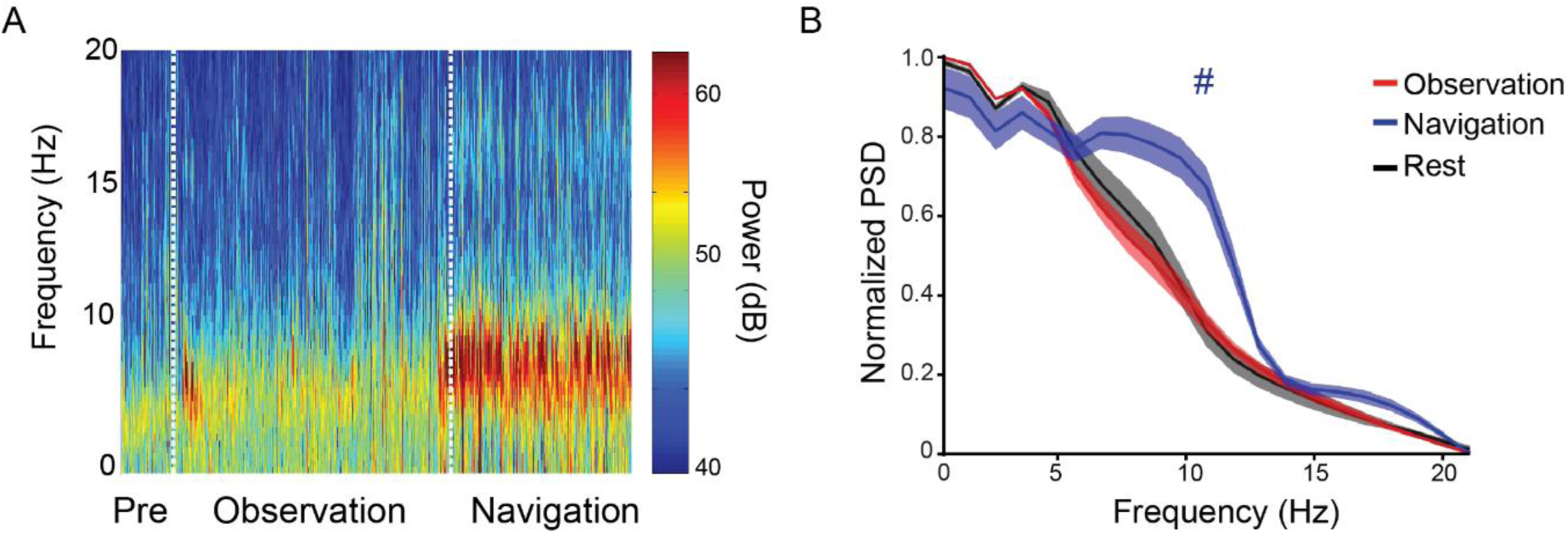
No changes in theta rhythm spectral power during observation. **(A)** Spectrogram (0-20 Hz) during a pre-task period of resting (Pre), observation and navigation phases of the learning task. Dotted vertical white lines indicate the transition between the different phases of the task. **(B)** Mean ± sem of power spectral density in the 0-20 Hz frequency range during the three phases of the task showed in (A). #p<0.05 navigation vs rest and vs observation, Friedman rank test followed by Dunn’s posthoc multiple comparisons.

## DISCUSSION

Observational learning is a relevant cognitive ability present in different species from invertebrate to human which able us to learn through the observation. Even though it has been demonstrated in different animal models at the behavioral level and some brain structure has been involved such as the amygdala, cerebellum, and hippocampus. The causal relationship between that structure and the capacity to learn by observation has been poorly explored. Recent evidence in rats and bats indicates that the principal cells of the observer hippocampus may encode the spatial position of another congener acting as a demonstrator. We hypothesized that the hippocampus is a critical structure for observational spatial learning through the ability to generate cognitive maps representing the experience of others or self.

Our results demonstrate that naïve rats can effectively improve their learning performance if before solving a spatial task they observe an expert demonstrator conducting the same task. This improvement in performance cannot be explained by other factors such as habituation to the observation platform or seeing a congener randomly moving around the arena. Importantly, this observational learning phenomenon was only present if the demonstrator corresponded to an expert rat solving the maze. Previous studies have demonstrated that the cerebellum is important for associative motor learning mediated by social demonstrations (Leggio et al 2000). Here we focused on hippocampus because it is the most relevant structure for spatial learning (Schenk & Morris,1985). Our experiment of hippocampal inactivation during observation clearly indicates that observational learning requires the proper functioning of this structure to be behaviorally unfolded. Since it is difficult to deal with remnant effect of drug in the brain our reversible inactivation of the hippocampus fully recovers the functionality of this region since expert animals which received the same procedures of inactivation display a correct spatial learning indicating that any effect of the drug during the observation did not affect the functionality of this structure during the execution of the task, causally demonstrating the importance of hippocampal neural activity in this phenomena.

In terms of electrical properties of hippocampal principal cells, we corroborate that these neurons effectively displayed the content of spatial information higher than chance and generated a spatial map that can represent both the position of others and themselves in the arena. The 19.3 % of pyramidal cells were classified as social place cells and 46.8% of social place cells also code for the animal’s own position during navigation (common cells). This percentage of social place cells is lower compared with Denjo et al (2018) study. This difference can be explained by the different methods used to define a place cell (presence of a place field vs. spatial information index) and by experimental design, e.g. in this present study observer animals were totally naïve respect to the task, and only observed without navigation during observation, while both in Denjo et al and Omer et al, observer animals must be trained to solve that particular task.

This difference in social place cells activated in experienced vs naïve animals, in turn, suggests that the experience of the animal before the observational learning experience could be critical for the hippocampal internal representation of other’s positions. This explanation could be extended to Mou & Ji et al., 2016 report, who did not find social place cell activity but report coactivation of place cells during observation, in an animal that passively watched a demonstrator in a linear track.

In terms of single-cell properties, social place cells share many features with canonical place cells, including average firing rate, spatial information, and sparsity. Only the firing rate at the center of the place field was higher during navigation respect to observation suggesting a remapping phenomenon between the 2 phases of the task. The neuronal population analysis demonstrates that the representation during navigation and observation is different with changes in firing rate overlap, global remapping and population vector representation of the space. These results indicate that the neuronal representation of what it is observed and explored are totally different, and the brain can differentiate between the experience of a congener and the self-animal but using the same neurophysiological substrate for both representations. How the observer’s brain manages the information of the demonstrator’s spatial performance in the generation of the self-spatial map, and how exactly, the non-spatial information contained in the social place cells drives observational learning, are questions that arise from our results and need further directions.

During navigation, the place cell activity is phase-locked with theta oscillation (Colgin, 2016). This coordinated pattern of activity has demonstrated to be necessary for spatial learning and it is proposed as a mechanism that may facilitate the transfer of information from other hippocampal formation structures (McNaughton et al. 2006). An increase in theta rhythm-power has been recently reported in social place cells (Danjo et al., 2018). However, we did not find any increase in theta power during the observation phase in our task. Given that place fields representation could emerges in absence of theta oscillation (Brandon et al., 2014) and ambulatory signals (e.g. quite animals; Terrazas et al., 2005), and the main difference between Danjo et al. (2018) and the present work is the degree of training of animals in the task, previous to observation (which in our case is not existent, because observers are naïve respect to the spatial task), and the animal is not navigating the maze during the observation, we suggest that theta power increase during observation could dependent on previous experience of the animal with the environment, but is not essential for the generation of a spatial representation of the demonstrator.

In summary, our report demonstrated that spatial observational learning depends on hippocampus, this structure can code the spatial representation of others and itself and that this coding is essential to learn from other’s experience.

## ACKNOWLEDGMENTS

We thank Dr. M. Schwalm and Dr(c) M. Toro-Espejo for their constructive comments of the initial versions of this manuscript. This work was supported by ICM P09-015F Biomedical Neuroscience Millennium Institute, ICM P-10-001-F Center for the memory neuroscience Millennium Nucleus, Science and Technology National Found grant # 11090294 and National Commission for Science and Technology Doctoral fellow #21150519.

## SUPPLEMENTARY FIGURES

**Figure S1.**
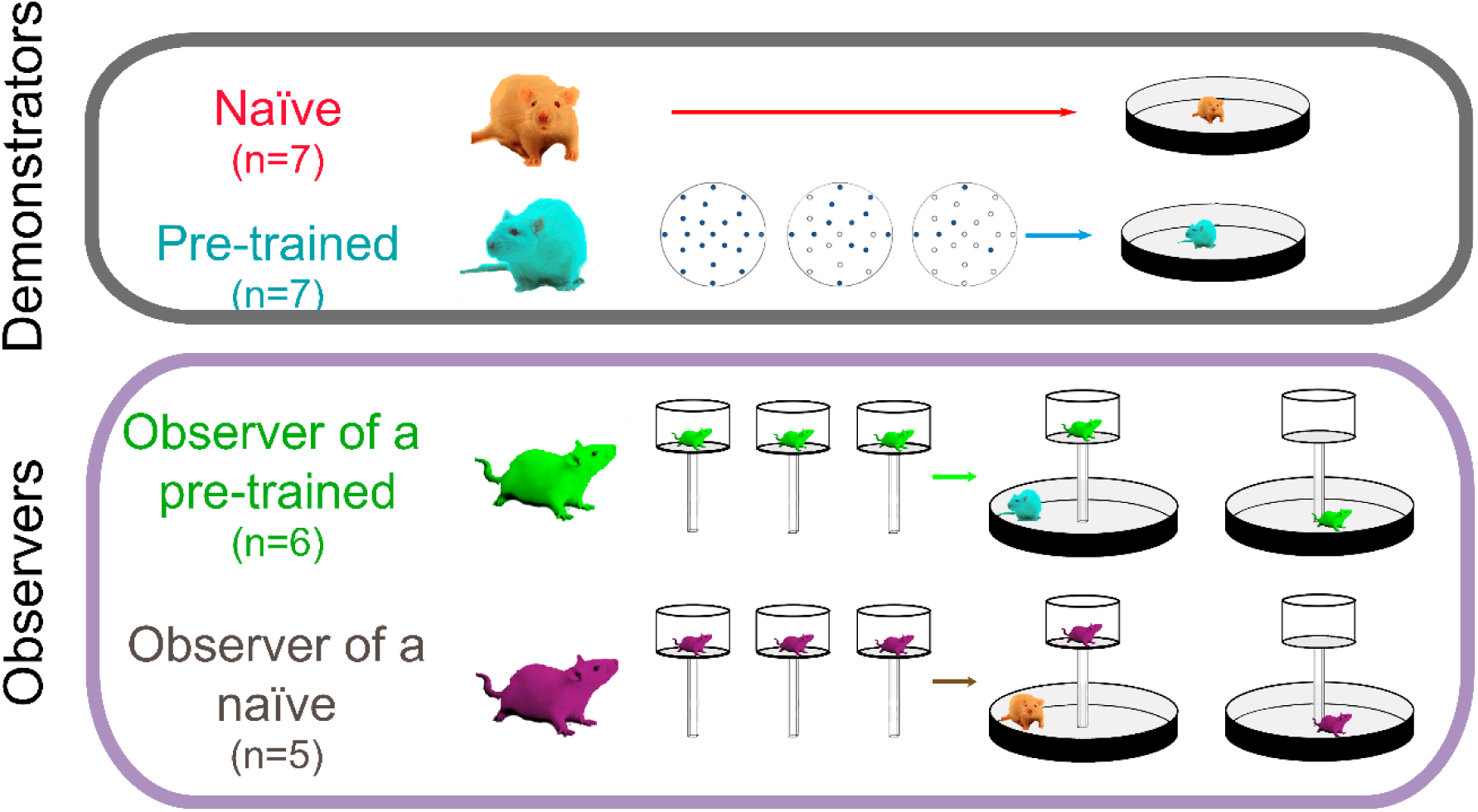
Behavioral groups and training details. A total of 25 rats were grouped in Demonstrators and Observers. Demonstrated were separated in naïve (red, n=7) and pre-trained demonstrators (blue, n=7). Observers were all naïve animals and grouped accordingly to the demonstrator nature. An observer of a pre-trained demonstrator (green, n=6) or an observer of a naïve (purple, n=5). The pre-trained protocol consisted of three consecutive days of free exploration of the oasis maze arena where 100, 50 and 25% of the wells were rewarded. Observer animals were habituated for three days to the observation platform and then watched the performance of a demonstrator in the oasis maze. Immediately after observation, those animals solved the oasis maze preserving the position of the baited well.

**Figure S2.**
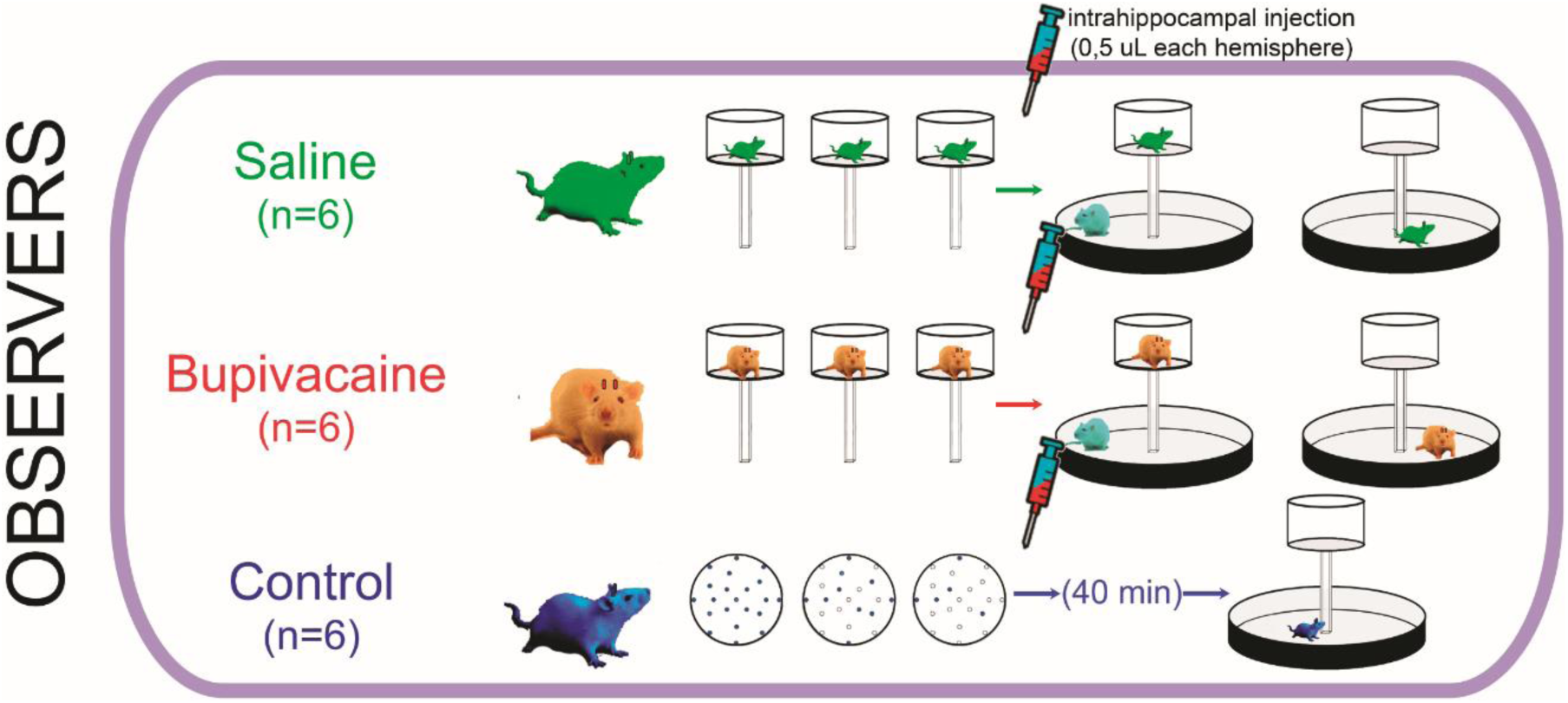
Pharmacological groups. A total of 18 animals were bilaterally implanted with cannulas directed to the hippocampal CA1 region. Six animals were injected with saline before the observation phase of the task (saline, green group), while the other 6 were bilaterally injected with 0.5 uL of bupivacaine 0.75% v/v (bupivacaine, red group). These two groups were previously habituated to the observational platform in three consecutive days. The last 6 animals were pre-trained in the oasis maze and 40 minutes before solving the task were injected with 0.5 uL of bupivacaine 0.75 % v/v, to mimic the time of observation phase and evaluate any residual effect of the drug in spatial navigation (control, blue group).

**Figure S3.**
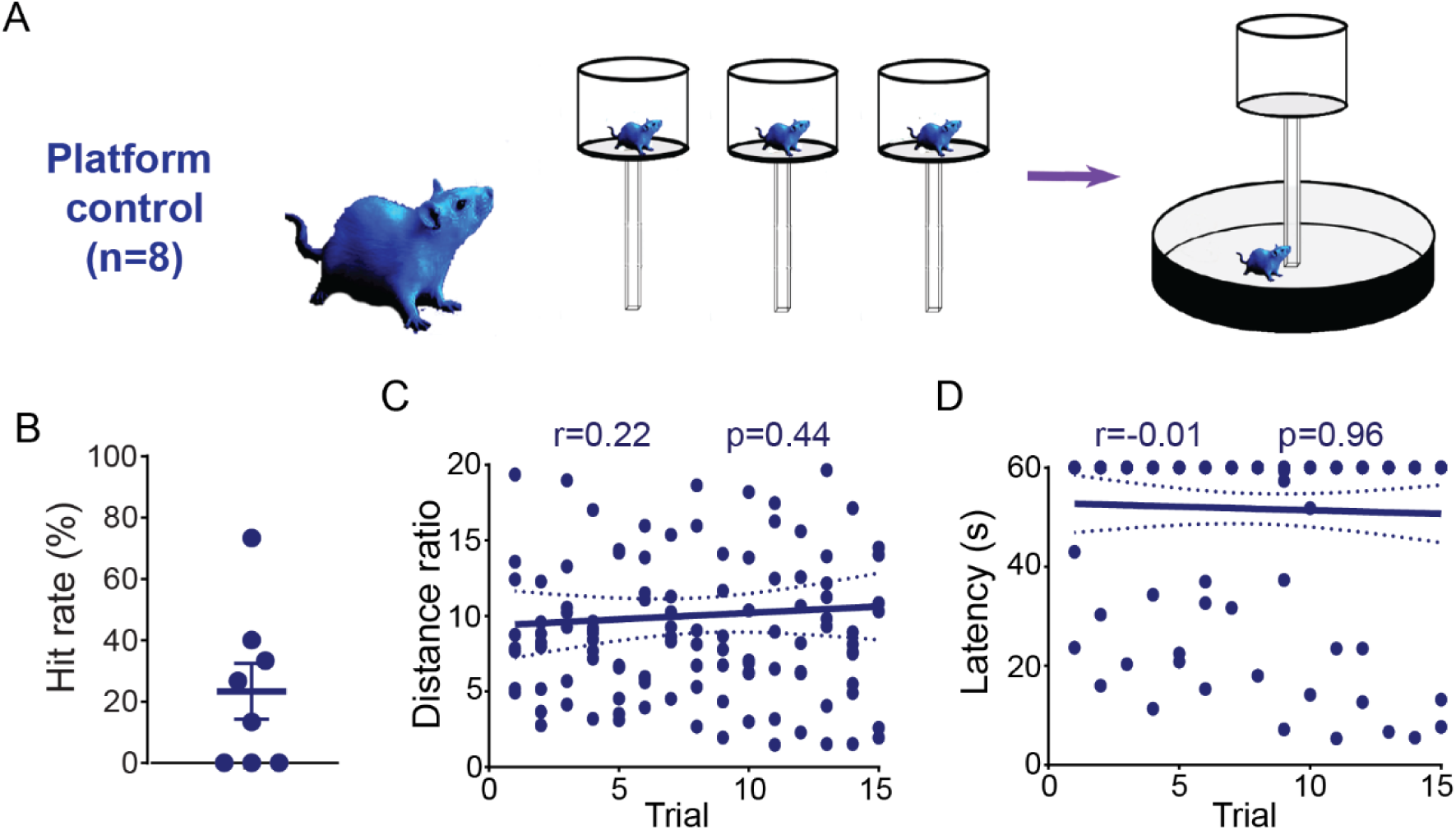
Platform control animals. **(A)** A subgroup of naïve rats (n=8) was habituated during three consecutive days to the platform and then solved the oasis maze without observing any demonstrator. **(B)** Mean ± sem percentage of hit rate, **(C)** distance ratio, **(D)** latency, the correlation coefficient and p-value of each behavioral variables it is indicated in the plot (Pearson correlation coefficient).

**Figure S4.**
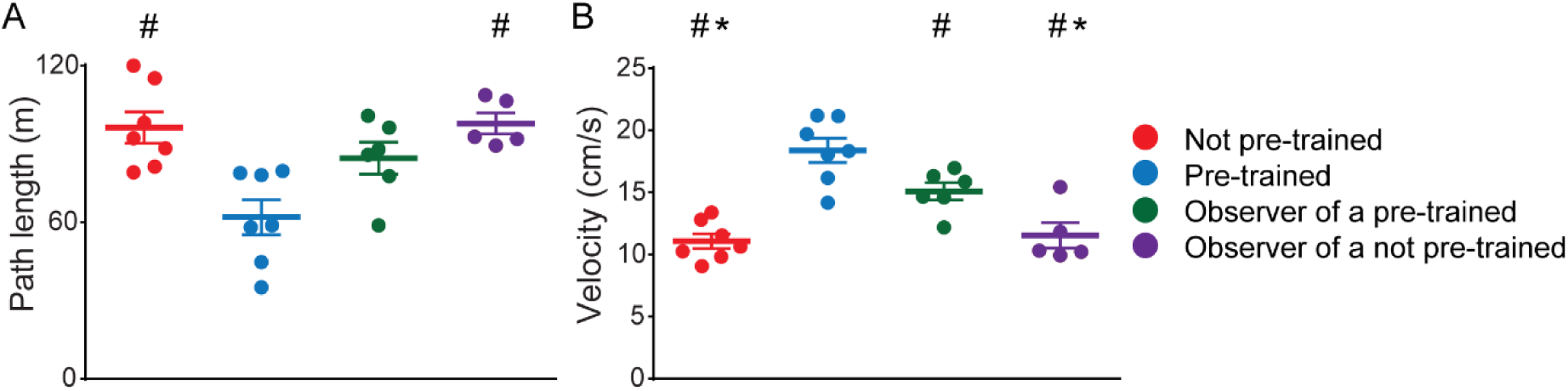
Poor spatial learning performance is not explained by lower exploration or immobility. Total path length **(A)** and mean velocity **(B)** during oasis maze. Results are shown as mean±sem. #p<0.05 vs pre-trained group, *p<0.05 vs observer of a pre-trained demonstrator, one-way ANOVA followed by Holm-Sidak posthoc test (for path length) and Kruskal-Wallis test followed by Dunn’s posthoc for velocity. Notice that the worst performance correspond to the animals that most distances traveled in the oasis maze.

**Figure S5.**
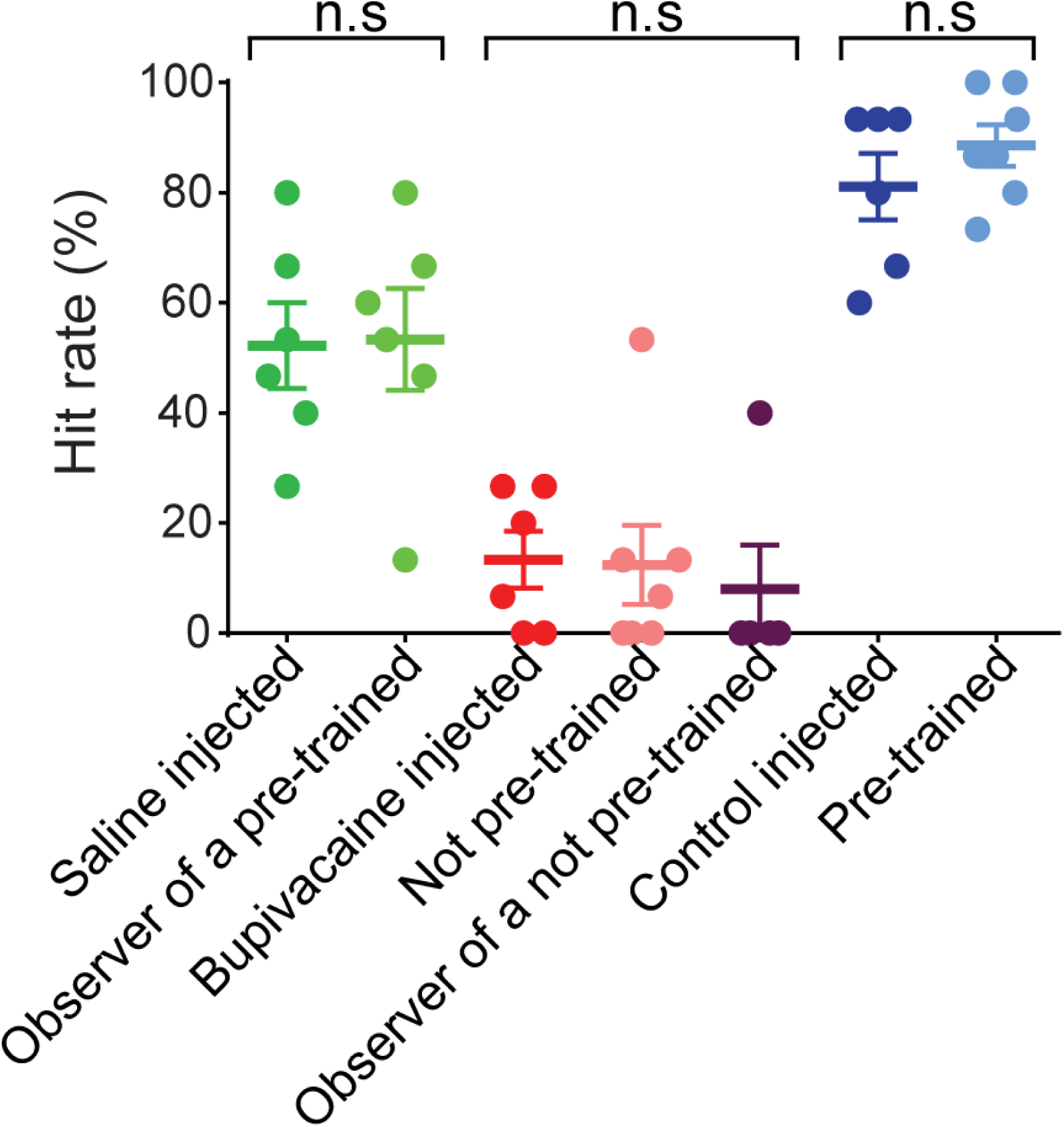
Comparison between behavioral and pharmacological hit rate. Mean ± sem of saline (green), an observer of a pre-trained (light green), bupivacaine (red), naïve demonstrator (pink), an observer of a naïve (purple), control (blue) and pre-trained (light blue) groups. n.s = non-significative, t-test (green vs light green and blue vs light blue groups) or Kruskal-Wallis one-way ANOVA (red, pink and purple groups).

**Figure S6.**
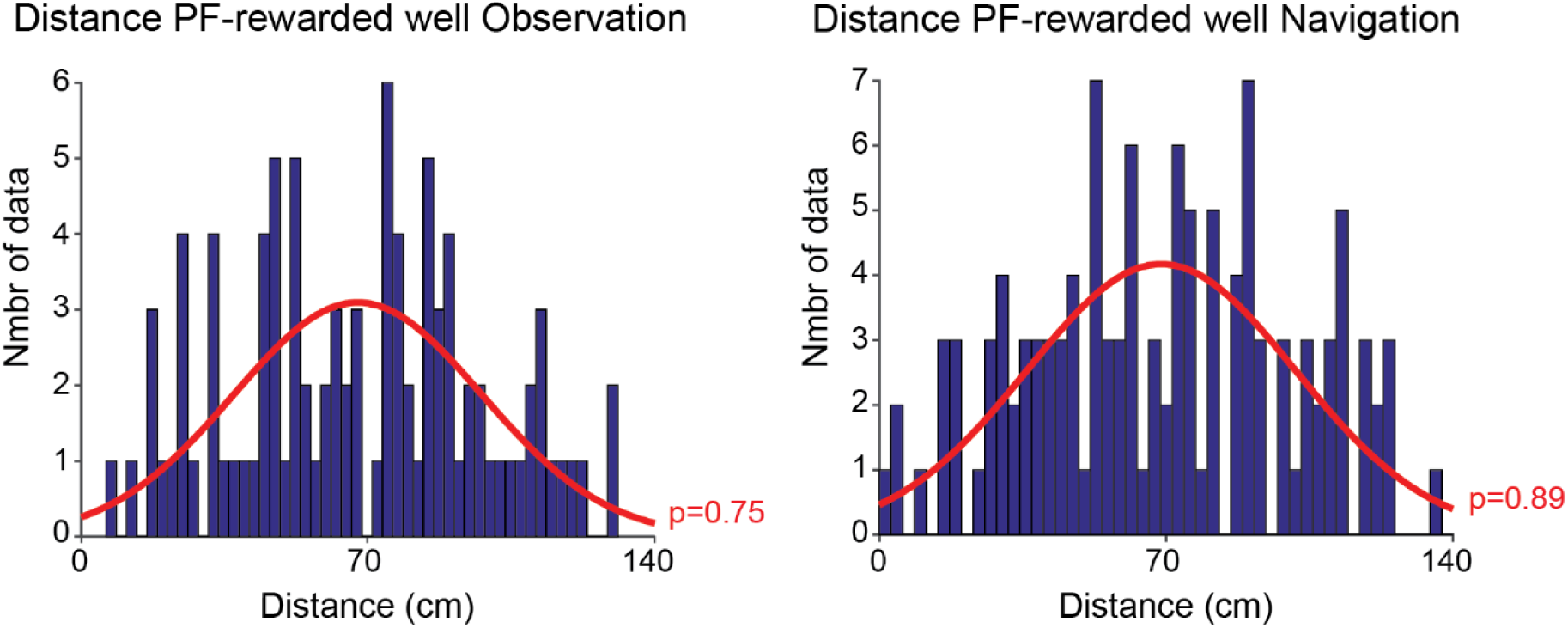
Place fields are not influenced by the position of the rewarded well. Distribution histogram for the distance between the center of the place field and the position of the rewarded-well both during observation (left) and navigation (right). The red curve represents a Gaussian distribution fitted with the distance values (p-value of a Kolmogorov test is indicated in red).

